# Assessing macrophyte seasonal dynamics using dense time series of medium resolution satellite data

**DOI:** 10.1101/279448

**Authors:** Paolo Villa, Monica Pinardi, Rossano Bolpagni, Jean-Marc Gillier, Peggy Zinke, Florin Nedelcuţ, Mariano Bresciani

**Affiliations:** Institute for Electromagnetic Sensing of the Environment, National Research Council, Milan, Italy; Department of Chemistry, Life Sciences and Environmental Sustainability, University of Parma, Parma, Italy; SNPN, Réserve naturelle du lac de Grand-Lieu, Bouaye, France; Norwegian University of Science and Technology, Trondheim, Norway; Dunărea de Jos University of Galati, Brăila, Romania

**Keywords:** Vegetation phenology, LAI, Shallow lakes, Spectral Indices, Sentinel-2, Landsat 8

## Abstract

Thanks to the improved spatial and temporal resolution of new generation Earth Observation missions, such as Landsat 8 and Sentinel-2, the potential of remote sensing techniques in mapping land surface phenology of terrestrial biomes can now be tested in inland water systems.

We assessed the capabilities of dense time series of medium resolution satellite data to deliver quantitative information about macrophyte phenology metrics, focusing on three temperate European shallow lakes with connected wetlands, located in Italy, France and Romania.

Leaf area index (LAI) maps for floating and emergent macrophyte growth forms were derived from semi-empirical regression modelling based on the best performing spectral index, with an error level around 0.11 m^2^ m^-2^. Phenology metrics were computed from LAI time series using TIMESAT code and used to analyse macrophyte seasonal dynamics in terms of spatial patterns and species-dependent variability. Peculiar patterns of autochthonous and allochthonous species seasonality across the three study areas were related to the environmental characteristics of each area in terms of ecological and hydrological conditions.

In addition, the influence of satellite dataset characteristics – i.e. cloud cover thresholding, temporal resolution and missing acquisitions – on phenology timing metrics retrieval was assessed. Results have shown that with full resolution (5-day revisit) time series, cloud cover can bias phenology timing metrics by less than 2 days, and that reducing temporal resolution to 15 days (similar to Landsat revisit) still allows for mapping the start and peak of macrophyte growth with an error level around 2–3 days.

## 1. Introduction

Although much is known about the long-term effect of climate change on the phenology of terrestrial ecosystems (Richardson et al., 2013; Yang et al., 2015), there is little information regarding aquatic ecosystems and even less regarding aquatic plants. Improving the knowledge on macrophyte seasonal changes or phenology is essential for investigating ecological drivers of aquatic systems degradation and promoting effective conservation programs. Some studies exist that focus on emergent macrophytes (Alahuhta et al., 2011), floating plants (Peeters et al., 2013) and their interaction with submerged macrophytes (Netten et al., 2011; Li et al., 2017). These works are mostly based on existing thematic cartography and *in situ* observations of vegetation density and biomass, thus not covering spatial and temporal scales that allow transferable if not general conclusions. Large lakes and wetland ecosystems are in fact difficult to survey and consequently few data have been collected to describe temporal dynamics of aquatic vegetation, as stressed recently by Luo et al. (2016). Consistent and spatialized data about key phenological features, e.g. the timing of start and end of growing season, are required to better understand the main drivers of aquatic vegetation seasonal dynamics (Wang et al., 2012; Sletvold and Ågren, 2015). Punctual knowledge of seasonal dynamics is also crucial for understanding the arrangement in space and time of macrophyte communities and functional groups, competition with other primary producers (Bolpagni et al., 2014; Villa et al., 2015; Zhang et al., 2015), as well as potential impact of invasive species (Wolkovich and Cleland, 2011). Furthermore, natural resource managers and policy makers demand knowledge of phenological dynamics over increasingly large temporal and spatial extents for addressing important issues related to global environmental change (White and Nemani, 2003; Cleland et al., 2007).

Within this frame, remote sensing offers near ideal capabilities in terms of spatial and temporal resolution, synoptic view and coverage, sensitivity to vegetation features (structural and physiological), regular sampling and operational repeatability (Malthus, 2017). Long-term, consistent satellite data are needed to monitor and quantify intra- and inter-annual trends in vegetation change (e.g. Villa et al., 2012; Fensholt et al., 2015). A large corpus of scientific literature on remote sensing of land surface phenology has been built in the last decade, focusing in particular on terrestrial biomes (e.g. Reed et al., 1994; Zhang et al., 2003; Fisher and Mustard, 2007). The majority of these works use satellite data with medium to low resolution, i.e. greater than 1 km pixel size (e.g. Jenkins et al., 2002; Reed, 2006; Fisher et al., 2007). Because of the peculiar characteristics of macrophytes (e.g. background conditions, canopy structures) and of their ecosystems (mostly shallow water bodies, with small surfaces and high heterogeneity in species and growth forms), techniques designed for making use of coarse spatial resolution data and targeted at terrestrial plants are not directly applicable and need to be adapted and/or reparametrized. Some studies for the phenological analysis of macrophytes were made using the 30m spatial resolution of the Landsat constellation sensors (e.g. Hestir et al., 2015; Luo et al., 2016), but the 16-day revisiting time does not ensure full coverage of macrophytes growth temporal variability.

Sentinel-2 satellite constellation, managed by ESA through the Copernicus initiative, started in July 2017 to provide high quality data at 10-20 m resolution, every 5 days (Drusch et al., 2012). Sentinel-2 data, for their characteristics, can be a powerful tool for monitoring macrophytes and their seasonal dynamics with a level of detail never experienced before.

In this paper, we analyse the capabilities of dense time series (5 to 10 day revisit) of medium resolution (10 to 30 m) satellite data to deliver information about macrophyte phenology metrics. For this purpose, we collected data and ran the analysis over three European shallow lakes with connected wetlands, hosting macrophyte communities of floating and emergent species, common to temperate freshwater ecosystems.

In particular, our objectives were: i) to calibrate a semi-empirical model for deriving Leaf Area Index (LAI) time series for floating and emergent macrophytes from satellite spectral reflectance data; ii) to map macrophyte phenology metrics and assess their spatial and species-dependent variability across our study areas; iii) to evaluate the influence of satellite dataset characteristics – i.e. maximum cloud cover amount, temporal resolution, and missing acquisitions – on phenology timing metrics derived.

## 2. Study areas

The study areas were three shallow freshwater bodies with connected wetlands, hosting macrophyte communities mainly composed by floating and emergent species. The three areas share a temperate climate and are located in Europe: Mantua lakes system (Italy) is the main study area, where extensive *in situ* data collection was carried out for implementing our analysis; Lac de Grand-Lieu (France), and Fundu Mare Island (Romania) are two additional test site areas, for which less reference information were available. Table 1 summarizes the key characteristics of main macrophyte species (floating and emergent) present in the study areas.

**Table 1.** Key characteristics of main target macrophytes species present in the study areas.

| Study area | Species<br>(common name) | Functional<br>group<sup>1</sup> | Abundance | Species origin |
| --- | --- | --- | --- | --- |
| Mantua lakes system | *Nelumbo nucifera*<br>(sacred lotus) | Emergent<br>rhizophyte | dominant<br>(Superior Lake) | allochthonous |
|  | *Trapa natans*<br>(water chestnut) | Free-floating<br>pleustophyte | dominant<br>(Middle and Inferior lakes) | autochthonous |
|  | *Nuphar lutea*<br>(yellow water lily) | Floating-leaved<br>rhizophyte | spread patches<br>(all lakes) | autochthonous |
|  | *Nymphaea alba*<br>(white water lily) | Floating-leaved<br>rhizophyte | small patches<br>(Middle Lake) | autochthonous |
|  | *Nymphoides peltata*<br>(yellow floating heart) | Floating-leaved<br>rhizophyte | small patches<br>(Vallazza wetland) | autochthonous |
|  | *Ludwigia hexapetala*<br>(water primrose) | Floating<br>rhizophyte | spreading<br>(Superior and Middle lakes) | allochthonous |
| Lac de Grand Lieu | *Nuphar lutea*<br>(yellow water lily) | Floating-leaved<br>rhizophyte | co-dominant | autochthonous |
|  | *Nymphaea alba*<br>(white water lily) | Floating-leaved<br>rhizophyte | co-dominant | autochthonous |
|  | *Nymphoides peltata*<br>(yellow floating heart) | Floating-leaved<br>rhizophyte | small patches | autochthonous |
|  | *Trapa natans*<br>(water chestnut) | Free-floating<br>pleustophyte | small patches | autochthonous |
|  | *Ludwigia hexapetala*<br>(water primrose) | Floating<br>rhizophyte | spreading<br>(riparian areas) | allochthonous |
|  | *Ludwigia peploides*<br>(creeping water primrose) | Floating<br>rhizophyte | spreading<br>(riparian areas) | allochthonous |
| Fundu Mare island | *Nymphaea alba*<br>(white water lily) | Floating-leaved<br>rhizophyte | co-dominant | autochthonous |
|  | *Trapa natans*<br>(water chestnut) | Free-floating<br>pleustophyte | co-dominant | autochthonous |
|  | *Nymphoides peltata*<br>(yellow floating heart) | Floating-leaved<br>rhizophyte | small patches | autochthonous |

<sup>1</sup> according to Villa et al. (2015) scheme

### 2.1. Mantua lakes system

The Mantua lakes system is located in the Po river floodplain (northern Italy; 45°10’ N, 10°47’ E; Figure 1c) with a continental climatic regime (Peel et al., 2007). The Superior, Middle and Inferior lakes are semi-artificial lakes created by damming the Mincio River in the 12^th^ century. The three fluvial lakes are small (~ 6 km^2^), shallow (average depth of 3.5 m), and hypertrophic (chlorophyll-a concentrations up to 100 μg L^-1^). The Superior Lake maintain a constant water level (17.5 m a.s.l.) due to water discharge regulation by the Vasarone sluice gate and Vasarina gate (Pinardi et al., 2015). In the Middle and Inferior lakes, water level is varying in very narrow range (14.0-14.5 m a.s.l.), for hydraulic safety (e.g. avoid flooding in the historic city centre). The lakes are protected as Natural Regional Park and are surrounded by two wetlands (Valli del Mincio and Vallazza; VM and VW), which are Nature Reserves. The Mantua lakes system is characterized by the coexistence of phytoplankton and macrophyte communities (Pinardi et al., 2011; Bolpagni et al., 2014; Villa et al., 2015). During the vegetative period (April-October) dense stands of *Nelumbo nucifera*, an allochthonous emergent rhizophyte, colonize the Superior Lake, together with some small patches of autochthonous floating species, *Nuphar lutea* and *Trapa natans*. The Middle Lake hosts dominant monospecific stands of *T. natans*, with little spots of nymphaeids (*N. lutea* and *Nymphaea alba*). The Inferior Lake mainly hosts small and isolated *T. natans* beds. In the last years, *Ludwigia hexapetala*, an allochthonous and highly invasive mat-forming species, has spread in the littoral zones of the Superior and Middle lakes.

**Figure 1.**
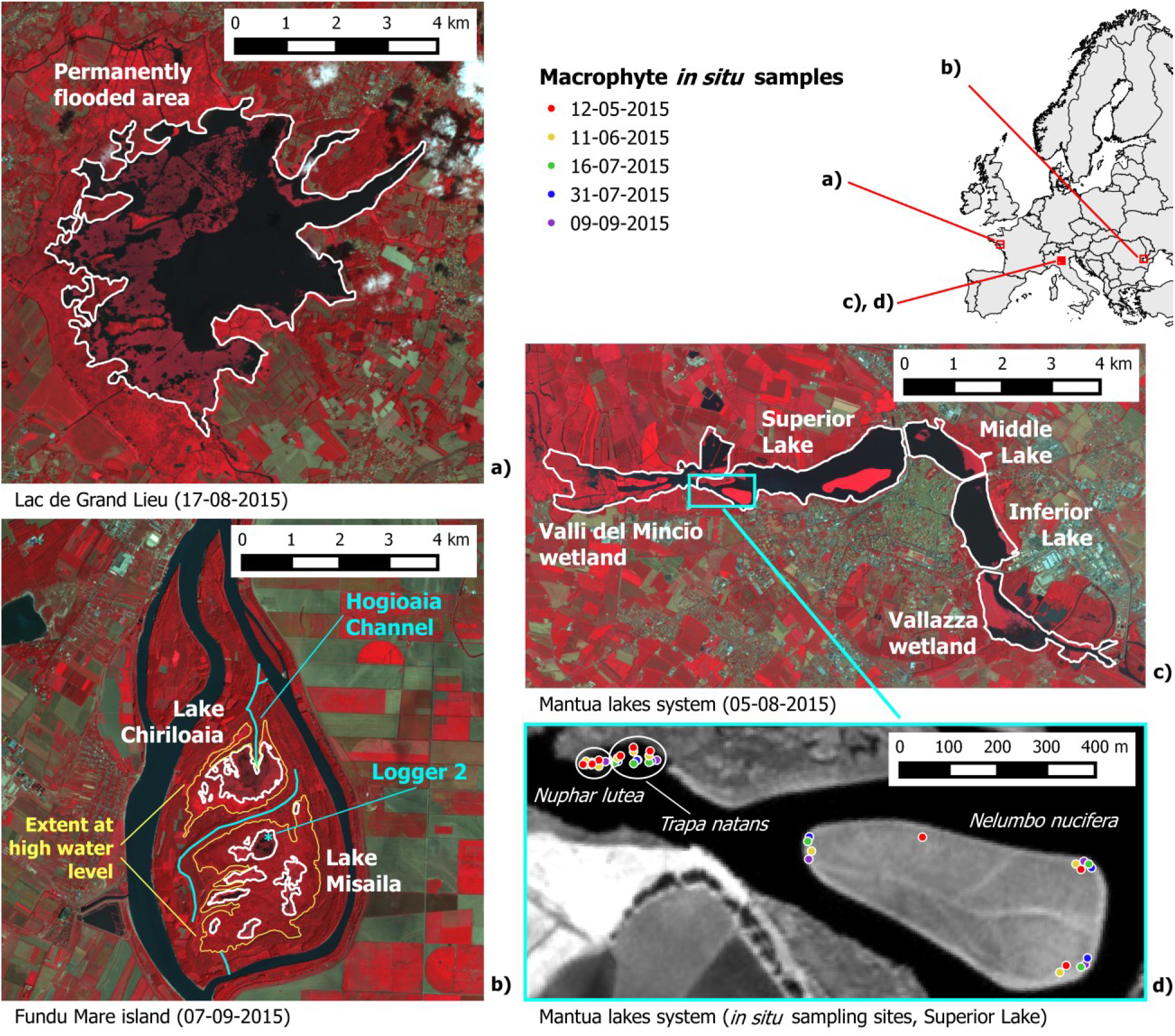
Study areas shown as SPOT5 images in colour infrared RGB combination at peak of macrophyte growth: a) Lac de Grand-Lieu (France); b) Fundu Mare Island (Romania); c) Mantua lakes system (Italy); d) detail area of Mantua lakes system with locations (see Table 1 for coordinates) of macrophyte samples collected *in situ* during 2015 (NIR band shown in greyscale).

### 2.2. Lac de Grand-Lieu

Lac de Grand-Lieu is a large, eutrophic and shallow freshwater lake in north-western France (Loire-Atlantique department, 47°05’ N, 1°41’ W; Figure 1a), located at 25 km from the Atlantic coast. It extends over 63 km^2^ during winter, when the wet meadows, reed beds, and tree groves (*Salix* spp., *Alnus* spp.) are flooded. Water level fluctuations follow the seasonal cycle of precipitations and artificial management. A sluice gate downstream of the lake controls water levels since the early 1960s. In spring, the water level drops by one meter (on average) with respect to the seasonal maximum. Then, during the summer, sluice gates are locked and water level slowly decreases according to the evaporation, until rainy autumn months. Depending on the year, the lake level may drop from 15 to 35 cm between July and October. The permanently flooded central area (21 km^2^), around one meter deep, is partially covered by floating-leaved macrophytes (~ 7 km^2^), dominated by *N. alba* and *N. lutea*. Little beds of *T. natans* and *Nymphoides peltata* extend over 0.1 to 0.3 km^2^ (Gillier and Reeber, 2016). Similarly to other nearby wetlands (Haury et al., 2011), the allochthonous, invasive *L. hexapetala* and *Ludwigia peploides* are widely spread in the channels and on the mudflats on the edge of the central area, expanding in a more “terrestrial” form on the wet meadows and on the sparse reed beds.

## 3. Dataset

### 3.1. In situ data over Mantua lakes system

*In situ* sampling campaigns were performed by boat during the vegetative period 2015 (12 May, 11June, 16 and 31 July, and 9 September) in Mantua lakes system, collecting samples of *N. lutea* (NL), *T. natans* (TN), and *N. nucifera* (NN) over a total of 45 plots (three species sampled in three sites, with replicates, in five dates). In each sampling location, plot represents an area homogeneously covered by a species, of around 10 m x 10 m (consistent with satellite data resolution).

For each macrophyte plot, *in situ* georeferenced (Trimble GeoXM) photos (RGB camera Sony DSC-HX60; three photos each plot) were taken from nadir, around 1 m above the canopy, framing a 1 m x 1 m floating square plot. Macrophyte LAI (m^2^ m^-2^) was derived by manually digitation of the areal size of each leaf (considering the overlapping of leaves) falling within the framed plot, for each image. As reported by Villa et al. (2017), this method is more accurate for floating and floating-leaved species than for species with emerging leaves (*N. nucifera*). As this work is focused on temporal vegetation evolution, a slight underestimation of LAI in absolute value for a single species is not considered limiting for the analysis of seasonal dynamics at ecosystem level, which is based on following the relative LAI curve along the growing season. Spectral response data of different surfaces, including terrestrial and aquatic targets, were acquired *in situ* for evaluating radiometric accuracy and consistency of processed satellite data (details in Supplementary Material).

### 3.2. Satellite data

Medium resolution (10–30 m ground pixel), broadband multi-spectral (visible to shortwave infrared wavelength range) satellite data were collected over the three areas, following the macrophyte seasonality of year 2015. The bulk of this dataset is composed by scenes acquired during the SPOT5 (Take5) experiment, during which SPOT5 data were collected over 150 sites every 5 days under fixed geometry from April to September 2015. SPOT5 (Take5) aimed to simulate the acquisition of time series that ESA’s Sentinel-2 full constellation started to provide with both its satellite operational, in July 2017. In order to complete the time series for covering the whole 2015, we used Landsat 7 and Landsat 8 scenes acquired earlier than April 2015, and Sentinel-2A scenes acquired after September 2015. The whole dataset (Figure 2) is composed of 33 scenes over the Mantua lakes system (plus 12 used for time series consistency checking), 27 scenes over the Lac de Grand-Lieu, and 29 scenes over the Fundu Mare Island.

**Figure 2.**
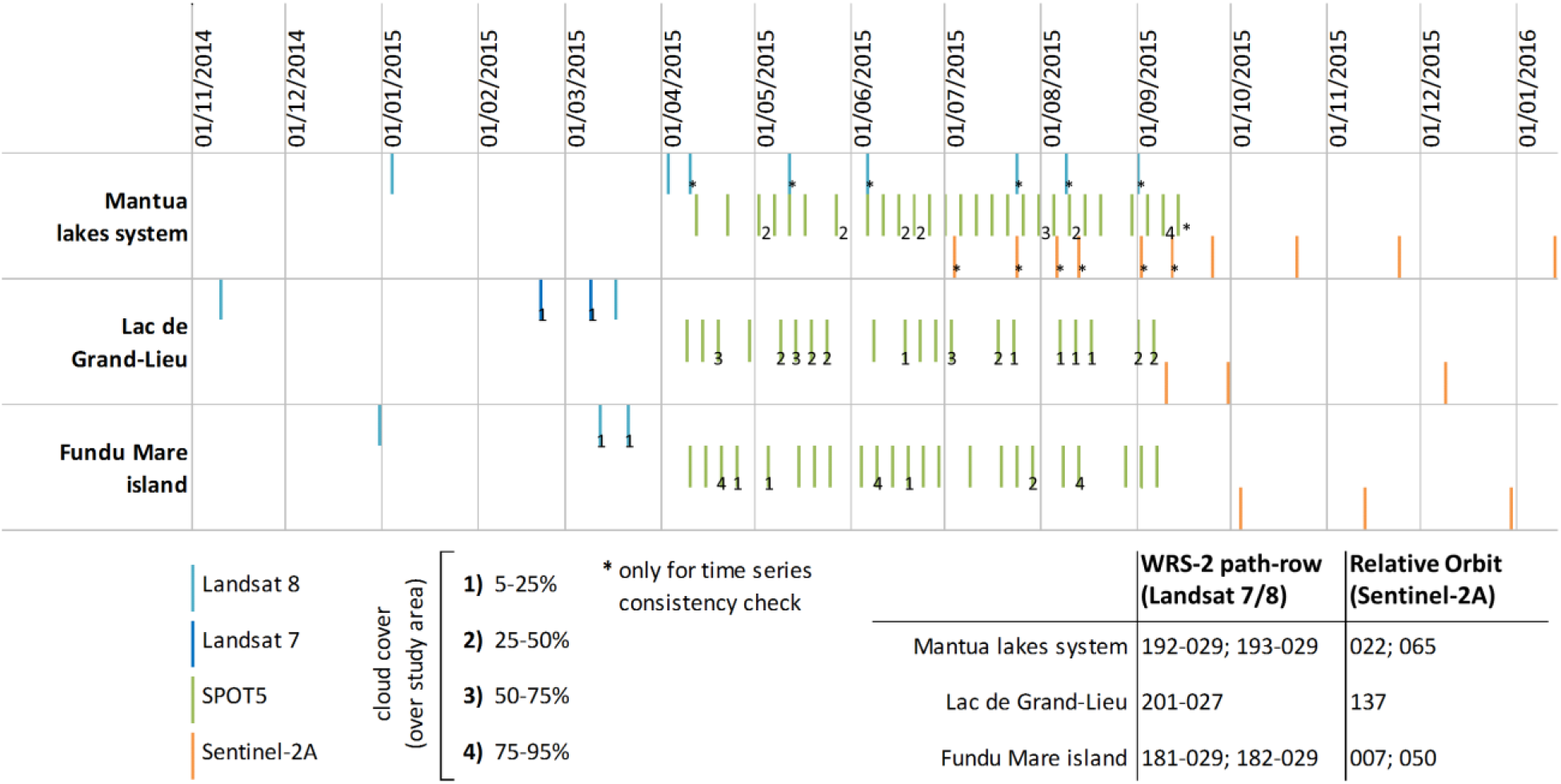
Satellite dataset characteristics.

### 3.3. Ancillary data

In Lac de Grand-Lieu, the monitoring of the floating-leaved plants is carried out every three years with the acquisition of high-resolution aerial RGB photos, covering the permanently flooded area and its surroundings (Figure 1a). In 2015, the survey took place on 8 August, with good weather conditions. For maintaining consistency with previous monitoring survey results, the 2015 orthorectified images were resampled at 0.5 m ground resolution and cropped to fit the area covered by floating and emergent macrophytes beds. In the resulting map, the main class merges *N. lutea* and *N. alba*, while other classes are *T. natans*, *N. peltata*, and *S. lacustris*. The areas covered by these four vegetation communities were digitized with a Geographic Information System (Quantum GIS).

Fundu Mare Island was visited on 8 May, 8 June and 21 December 2015, and a comprehensive field survey was carried out between 9 and 16 July 2015 in order to improve the understanding of the hydrological processes at the island. Between 9 June and 21 December 2015, the water level was recorded at three sites in the lakes using pressure sensors (Global Water). The water level upstream from the weir in the main outlet channel (Hogioaia channel, Figure 1b) was measured at four dates and its temporal development between these dates was estimated based on the water balance (Zinke et al. 2016). Figure 3 shows the water levels in 2015, with reference to the geodetic datum Black Sea Sulina (i.e. a.s.l.), and the typical water depths under different macrophyte communities characterizing the site.

**Figure 3.**
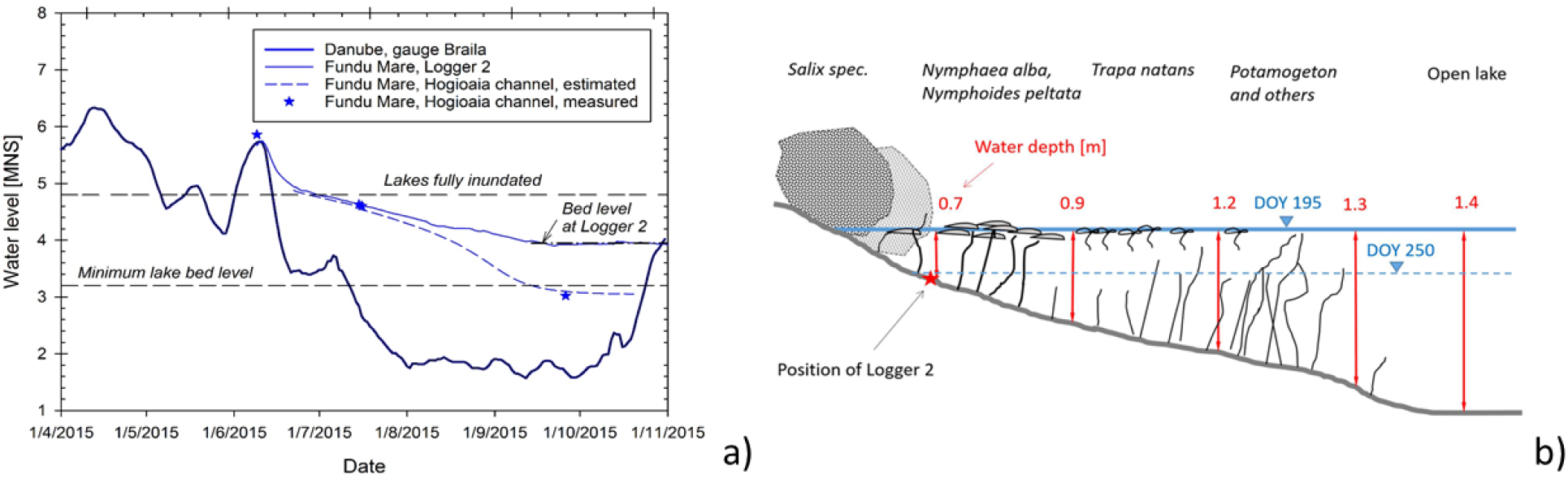
Relations between macrophyte and hydrology in Fundu Mare Island: a) water levels of the Danube (gauge in Braila city: 45°16’N, 27°57’E), in Lake Misaila (Logger 2, Figure 1b) and upstream from the weir (Hogioaia channel, Figure 1b). Note that 2015 winter season maximum level is higher than average levels of lakes when fully inundated (~4.8 m a.s.l.); during 2015 summer, the lake water level dropped below the bed elevation at the logger site (3.9 m a.s.l.); b) sketch of typical water depths measured under different macrophyte communities during the field survey on 14 July 2015 (DOY 195), at 4.6 m a.s.l.. Position of Logger 2 in Lake Misaila and water level on 7 September 2015 (DOY 250) are also shown.

The status of the vegetation during the field activities was documented by georeferenced photos taken with a RGB digital camera (Olympus, with GPS tagging), in combination with field notes about community types and species.

On 13 and 14 July 2015, water depth was measured in lakes Chiriloaia and Misaila (Figure 1b) at selected locations, ranging between 0.7 and 1.5 m (Figure 3b).

The overall vegetation cover in September 2015 was documented during an aerial survey performed with an Unmanned Aerial Vehicle (UAV; Parrot SenseFly eBee), mounted with a high-resolution digital camera (Canon IXUS 127 HS; 16.1 Mpixels). Roughly eight thousand photos were taken from around 200 m flight height, covering 20 km^2^, with nominal pixel size on the ground varying between 2 and 5 cm, that were used to generate an RGB orthomosaic of the whole area.

## 4. Methods

### 4.1. Satellite data pre-processing

SPOT5 data were retrieved from the ESA-CNES webportal (https://spot-take5.org), for the three sites that contain our study areas, namely: ‘Italia: Mantua’ (Mantua lakes system), ‘France: Pornic’ (Lac de Grand-Lieu), and ‘Romania: Braila’ (Fundu Mare Island). SPOT5 data were retrieved as Level 2A products: i.e. ortho-rectified surface reflectance, corrected from atmospheric effects (including adjacency) using the Multi-sensor Atmospheric Correction and Cloud Screening processor (MACCS; Hagolle et al., 2015).

Landsat scenes were pan-sharpened at 15 m resolution, using the Gram-Schmidt method (Laben and Brower, 2000). For Landsat 7 data, SLC-off gaps were filled using the approach developed by Maxwell et al. (2007).

Landsat and Sentinel-2A data were first radiometrically calibrated, and then converted to surface reflectance using ATCOR-2 code (Richter and Schläpfer, 2014). ATCOR-2 was run using image-based visibility estimation (Kaufman et al., 1997) and adjacency effect compensation (1 km radius).

Homologous spectral bands from the multi-sensor dataset were retained for further processing, by selecting the best matching bands covering the four broadband bands of SPOT5 data (Table 2).

**Table 2.** Homologous spectral bands used for building consistent reflectance time series per each sensor and satellite employed.

| Spectral range | Symbol | Corresponding spectral bands per each sensor (acronym, satellite) |  |  |  |
| --- | --- | --- | --- | --- | --- |
|  |  | Enhanced Thematic Mapper +<br>(ETM+, Landsat 7) | Operational Land Imager<br>(OLI, Landsat 8) | High Resolution Geometric inst.<br>(HRG, SPOT5) | Multi Spectral Instrument<br>(MSI, Sentinel-2A) |
| visible reflectance – green (Green) | $\rho_{Green}$ | Band 2 | Band 3 | Band 1 | Band 3 |
| visible reflectance – red (Red) | $\rho_{Red}$ | Band 3 | Band 4 | Band 2 | Band 4 |
| near-infrared reflectance (NIR) | $\rho_{NIR}$ | Band 4 | Band 5 | Band 3 | Band 8 |
| shortwave infrared reflectance (SWIR1) | $\rho_{SWIR1}$ | Band 5 | Band 7 | Band 4 | Band 11 |

Geometric and radiometric quality of multi-sensor satellite data were assessed, in order to check the dataset adequacy to common requirements for the inter-sensor geometric consistency and radiometric accuracy of spectral reflectance derived from different medium resolution platforms (SPOT5, Landsat 7/8, Sentinel-2A). Details are provided in Supplementary Material.

### 4.2. Modelling macrophyte LAI

Following the approach of Villa et al. (2017), macrophyte LAI maps were derived from satellite data through semi-empirical regression modelling based on spectral indices (SIs). Macrophyte LAI data collected *in situ* during 2015 growing season in Mantua lakes system, representing homogeneous macrophyte plots of about 10 m x 10 m, were split into two subsets (Table S1): two-thirds were used for calibrating (30 samples), and one third for validation (15 samples) of the semi-empirical LAI model implemented.

A range of ten SIs sensitive to vegetation canopy morphology and based on broadband surface reflectance in four spectral ranges (Green, Red, NIR, and SWIR1; see Table 2) matching SPOT5 bands were tested (Table 3, including related references). The 4 band broadband reflectance spectra of each macrophyte plot sampled in Mantua lakes system were derived from SPOT5 data (at 10 m pixel resolution) acquired within 5 days from *in situ* data collection. SPOT5 spectra were extracted from 3 x 3 pixels windows centred on the location of *in situ* samples, retaining the maximum vegetated pixel as matchup (Villa et al., 2017).

**Table 3.**
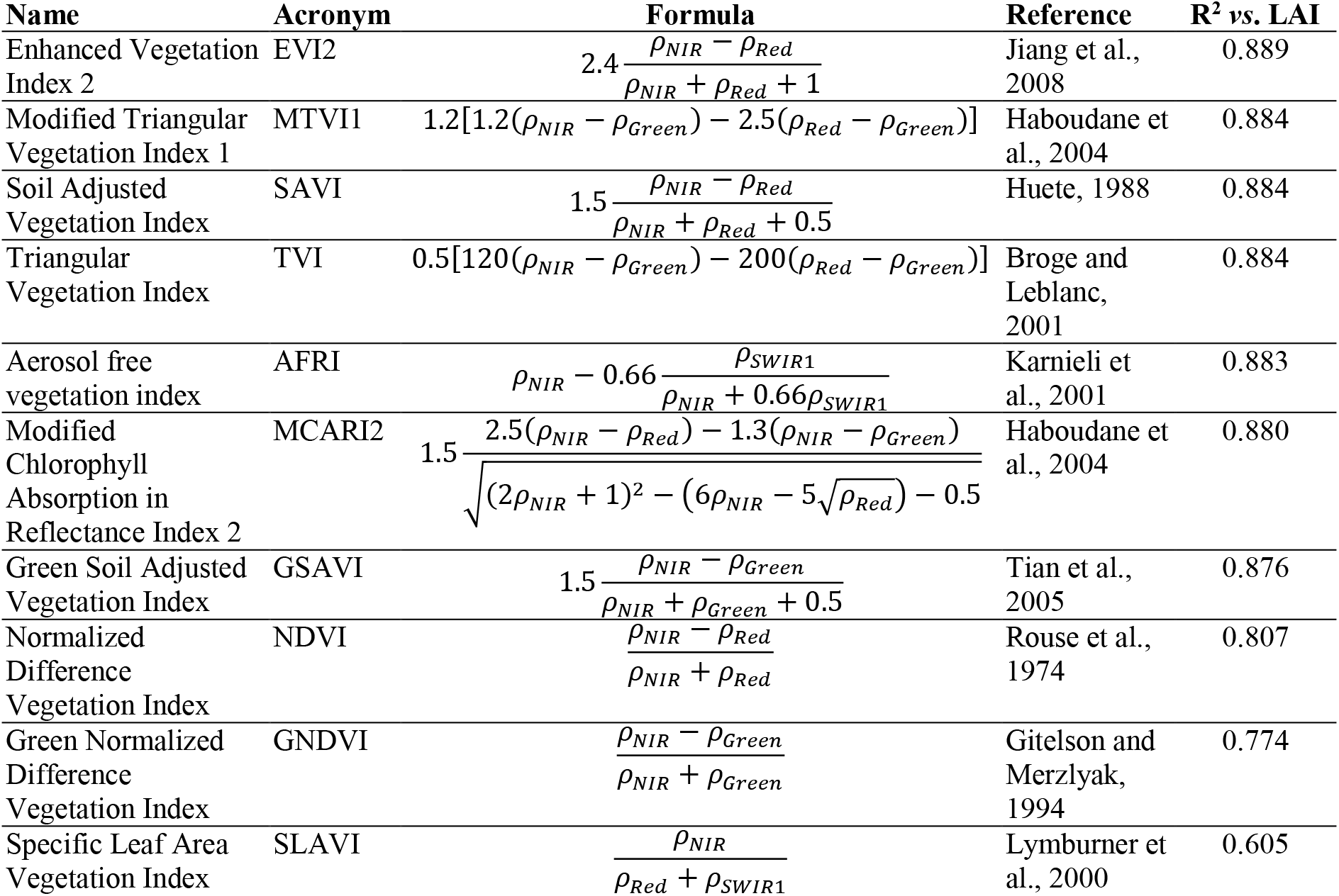
Spectral Indices tested, including index formulas, references and R^2^ with macrophyte LAI for the calibration set.

| Name | Acronym | Formula | Reference | $R^2$ vs. LAI |
| --- | --- | --- | --- | --- |
| Enhanced Vegetation Index 2 | EVI2 | $2.4 \frac{\rho_{NIR} - \rho_{Red}}{\rho_{NIR} + \rho_{Red} + 1}$ | Jiang et al., 2008 | 0.889 |
| Modified Triangular Vegetation Index 1 | MTVI1 | $1.2[1.2(\rho_{NIR} - \rho_{Green}) - 2.5(\rho_{Red} - \rho_{Green})]$ | Haboudane et al., 2004 | 0.884 |
| Soil Adjusted Vegetation Index | SAVI | $1.5 \frac{\rho_{NIR} - \rho_{Red}}{\rho_{NIR} + \rho_{Red} + 0.5}$ | Huete, 1988 | 0.884 |
| Triangular Vegetation Index | TVI | $0.5[120(\rho_{NIR} - \rho_{Green}) - 200(\rho_{Red} - \rho_{Green})]$ | Broge and Leblanc, 2001 | 0.884 |
| Aerosol free vegetation index | AFRI | $\rho_{NIR} - 0.66 \frac{\rho_{SWIR1}}{\rho_{NIR} + 0.66\rho_{SWIR1}}$ | Karnieli et al., 2001 | 0.883 |
| Modified Chlorophyll Absorption in Reflectance Index 2 | MCARI2 | $1.5 \frac{2.5(\rho_{NIR} - \rho_{Red}) - 1.3(\rho_{NIR} - \rho_{Green})}{\sqrt{(2\rho_{NIR} + 1)^2 - (6\rho_{NIR} - 5\sqrt{\rho_{Red}}) - 0.5}}$ | Haboudane et al., 2004 | 0.880 |
| Green Soil Adjusted Vegetation Index | GSAVI | $1.5 \frac{\rho_{NIR} - \rho_{Green}}{\rho_{NIR} + \rho_{Green} + 0.5}$ | Tian et al., 2005 | 0.876 |
| Normalized Difference Vegetation Index | NDVI | $\frac{\rho_{NIR} - \rho_{Red}}{\rho_{NIR} + \rho_{Red}}$ | Rouse et al., 1974 | 0.807 |
| Green Normalized Difference Vegetation Index | GNDVI | $\frac{\rho_{NIR} - \rho_{Green}}{\rho_{NIR} + \rho_{Green}}$ | Gitelson and Merzlyak, 1994 | 0.774 |
| Specific Leaf Area Vegetation Index | SLAVI | $\frac{\rho_{NIR}}{\rho_{Red} + \rho_{SWIR1}}$ | Lymburner et al., 2000 | 0.605 |

All matchup spectra (N=45) were then used to derive the SIs listed in Table 3, for both calibration and validation sample sets. The coefficient of determination (R^2^) between *in situ* macrophyte LAI and the SPOT5 derived SIs for the calibration set (N = 30; Table S1) was used as indicator of goodness of fit to inform the selection of the best SI for LAI modelling through linear regression (see last column of Table 3).

LAI model performance was assessed in terms of Mean Absolute Error (MAE), Mean Absolute Percentage Error (MAPE), and R^2^, calculated over the separate validation set (N = 15; Table S1). The macrophyte LAI model implemented was then applied to the satellite dataset covering our three study areas (Figure 2). LAI maps were produced only for floating and emergent macrophytes, which were isolated by using a binary raster mask (mask value = −1) composed by all pixels belonging to the water body area delineated using pre-season data (i.e. with maximum EVI2 < 0.05 in January-March), and covered by vegetation during the growing season (i.e. with maximum EVI2 > 0.1 in April-September).

Cloud covered areas, identified as pixels with *ρ_Green_* > 0.15 surrounded by 50 m buffer, were masked out from macrophyte LAI maps (mask value = −1). Subsequently, in order to homogenise spatial resolution of products derived from multiple sensors with different original resolution (Landsat, SPOT5 and Sentinel-2A), macrophyte LAI maps were all resampled at 20 m pixel size, using bilinear interpolation.

Time series of macrophyte LAI for the whole 2015 year with 5 day maximum temporal resolution – nominal revisit of SPOT5 Take5 dataset and Sentinel-2A/B joint constellation – were established by filling dates with missing satellite acquisitions with void layers (NA value = −1), i.e. for pre-April and post-September periods, covered respectively by Landsat and Sentinel-2A data (see Figure 2).

### 4.3. Metrics of seasonal dynamics

Quantitative descriptors of macrophyte seasonal dynamics were derived using TIMESAT software with macrophyte LAI maps as input (Jönsson and Eklundh, 2002; Jönsson and Eklundh, 2004). TIMESAT output parameters considered, hereafter called seasonal dynamics metrics, were (Figure 4): the time of the start of the season (SoS, in Day of Year: DOY), time for the peak of the season (PoS, in DOY), the time of the end of the season (EoS, in DOY), the maximum LAI value reached during the season (LAI_max, in m^2^_veg_ m^-2^), the rate of increase of LAI during the early vegetative phase (LAI_growth, in m^2^_veg_ m^-2^ d^-1^), the rate of decrease of LAI during the senescence phase (LAI_senescence, in m^2^_veg_ m^-2^ d^-1^).

**Figure 4.**
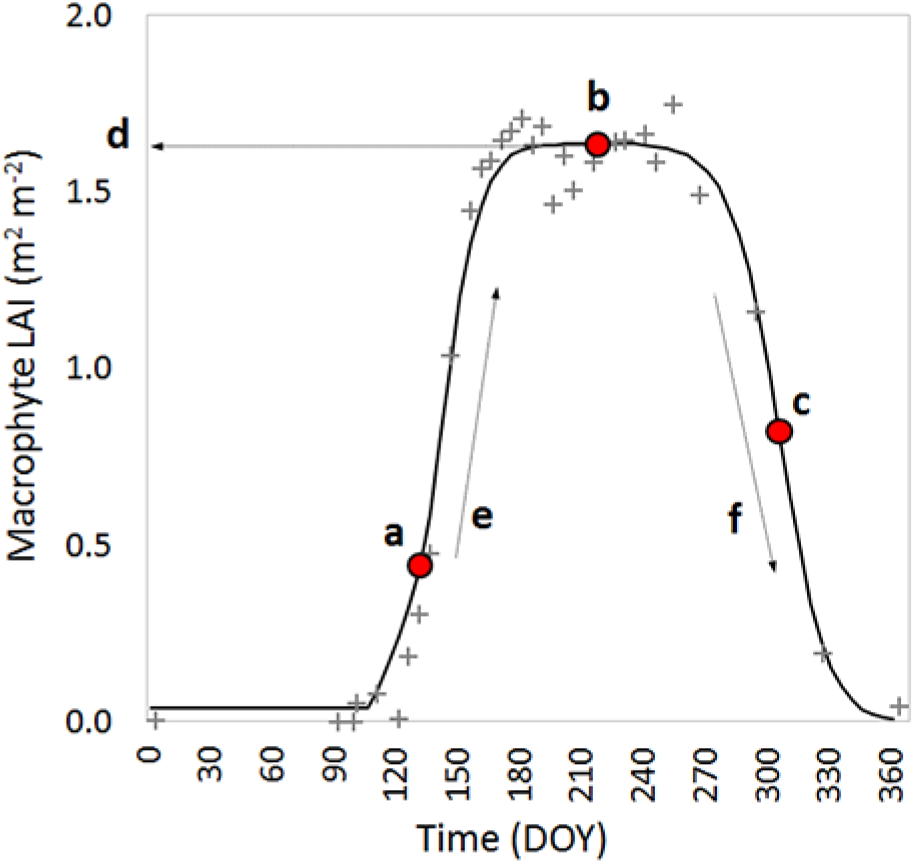
Metrics of seasonal dynamics derived from macrophyte LAI time series using TIMESAT: (a) SoS, (b) PoS, (c) EoS, (d) LAI_max, (e) LAI_growth, (f) LAI_senescence. The example is derived for a *Nelumbo nucifera* bed located in Mantua lakes system (45°09’40’’N, 10°46’30’’E). Grey crosses represent LAI values derived from satellite time series, and black line is the fitted asymmetric Gaussian curve. Adapted from Eklundh and Jönsson (2015).

TIMESAT was run without applying any spike filtering and setting the envelope iterations number to one. The curve fitting method selected was based on Asymmetric Gaussian curves, because it was demonstrated to be less sensitive to noise and incompleteness of input time series (Gao et al., 2008; Tan et al., 2011). According to our observations of macrophyte species investigated, the start of season occurs when macrophytes overpass 25% of peak LAI (i.e. 0.25 of the fitted curve amplitude, before PoS), and the end of season is flagged when macrophyte LAI decrease during senescence phase under 50% of reached maximum (i.e. 0.50 of the fitted curve amplitude, after PoS). For further details on TIMESAT requirements, capabilities and outputs the reader is referred to the software manual (Eklundh and Jönsson, 2015).

### 4.4. Influence of input variables on macrophyte phenology estimation

We evaluated the influence of some characteristics of the time series input dataset on the macrophyte seasonal dynamics results by assessing the variability of key seasonality metrics, namely the SoS, PoS, and EoS, when the input multi-temporal dataset is modified. In particular, we investigated the sensitivity of SoS, PoS, and EoS to i) could cover amount in input time series; ii) temporal resolution of input time series; and iii) relative importance of single dates or periods.

We carried out this analysis using the Mantua lakes system dataset, for which we collected field observations during 2015 growing season. We compared the SoS, PoS, and EoS derived from the baseline macrophyte LAI time series input, i.e. the time series with nominal time revisit of 5 days acquired during the SPOT5 Take5 experiment (hereafter named Mantua_5d, see Table 4), with the same seasonal metrics calculated when input to TIMESAT was varied as described in the following. The comparison was carried out by calculating differences in SoS, PoS, and EoS outputs over four selected macrophyte beds with different dominating species, environmental conditions and seasonality features (Figure S3). For this purpose, we calculated the Mean Absolute Difference (MAD), and the span between minimum and maximum values (Max_span) of SoS, PoS, and EoS derived from modified and baseline input dataset.

**Table 4.** Characteristics of LAI time series datasets used for investigating influence of input variables on estimating macrophyte phenology timing metrics.

| Target | Dataset name | Number of scenes | Short description |
| --- | --- | --- | --- |
| Baseline dataset | Mantua_5d | 33 | complete LAI time series at 5 day revisit |
| Testing influence of cloud cover amount | Mantua_5d_CC50 | 31 | modified input LAI time series with 50% maximum cloud cover allowed |
|  | Mantua_5d_CC30 | 29 | modified LAI time series with 30% maximum cloud cover allowed |
|  | Mantua_5d_CC10 | 26 | modified LAI time series with 10% maximum cloud cover allowed |
| Testing influence of temporal resolution | Mantua_10d_a | 20 | modified LAI time series with 10 day revisit (starting from DOY 2) |
|  | Mantua_10d_b | 19 | modified LAI time series with 10 day revisit (starting from DOY 7) |
|  | Mantua_15d_a | 15 | modified LAI time series with 15 day revisit (starting from DOY 2) |
|  | Mantua_15d_b | 15 | modified LAI time series with 15 day revisit (starting from DOY 7) |
|  | Mantua_15d_c | 15 | modified LAI time series with 15 day revisit (starting from DOY 12) |
| Testing influence of missing acquisitions | Mantua_5d_[1-31] | 30 | modified LAI time series with 5 day revisit and single dates excluded: steps [1-31] in Mantua_5d_CC50 |
|  | Mantua_15d_b_[1-15] | 14 | modified LAI time series with 15 day revisit and single dates excluded: steps [1-15] in Mantua_15d_b |

#### 4.4.1. Influence of could cover amount

For assessing the influence of cloud cover amount, three modified input dataset were prepared, by setting different thresholds on maximum cloud cover amount for each scene (Figure 2), thus excluding the dates which in baseline dataset show cloud cover exceeding this threshold (Table 4): 50% (2 scenes removed; Mantua_5d_CC50); 30% (4 scenes removed; Mantua_5d_CC30); 10% (7 scenes removed; Mantua_5d_CC10).

#### 4.4.2. Influence of temporal resolution

For assessing the influence of temporal resolution, five modified input dataset were prepared at 10-day and 15-day revisit (Table 4), respectively aimed to simulate Sentinel-2A and Landsat series revisit (16 days). Two dataset with 10-day temporal resolution were produced by removing one every two dates in the baseline dataset, starting from DOY 2 (Mantua_10d_a), or from DOY 7 (Mantua_10d_b). Three dataset with 15-day temporal resolution were produced by removing two every three dates in the baseline dataset, starting from DOY 2 (Mantua_15d_a), from DOY 7 (Mantua_15d_b), or from DOY 12 (Mantua_15d_c).

#### 4.4.3. Influence of missing acquisitions

For simulating the influence of missing dates within a yearly time series, possibly occurring because of cloud cover, extreme atmospheric conditions, or other issues due to the sensor (e.g. maintenance or failed acquisitions), 46 modified input dataset were prepared at 5-day and 15-day revisit (Table 4). 31 dataset with nominal 5-day temporal resolution were prepared, each produced by removing one single date in the LAI time series with 50% maximum cloud cover allowed (named Mantua_5d_[1-31], depending on which date is removed). 15 dataset with nominal 15-day temporal resolution were prepared, each produced by removing one single date in resampled LAI time series ‘Mantua_15d_b’ (named Mantua_15d_b_[1-15], depending on which date is removed).

## 5. Results

### 5.1. Modelling macrophyte LAI

EVI2, being the SI derived from SPOT5 scoring highest R^2^ with *in situ* macrophyte LAI for the calibration set (R^2^ = 0.889, Table 3), was used for implementing the semi-empirical LAI model through linear regression, using equation (1):

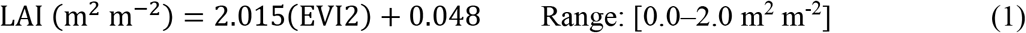

Comparison between LAI measured *in situ* and LAI model outputs (Figure 5) shows an overall error level of 0.11 m^2^ m^-2^ in absolute terms (MAE), for both calibration and validation sample sets, and lower than 18% in relative terms (MAPE of 17.7% and 9.6% for calibration and validation sets, respectively). Over both sample sets, the model tends to slightly underestimate LAI values higher than 1.5–1.6 m^2^ m^-2^ (corresponding to mature *N. nucifera* plots), with some mismatch at intermediate LAI values for *T. natans* (~20% overestimation of LAI > 1 m^2^ m^-2^) and *N. lutea* (15–25% underestimation of low LAI conditions).

**Figure 5.**
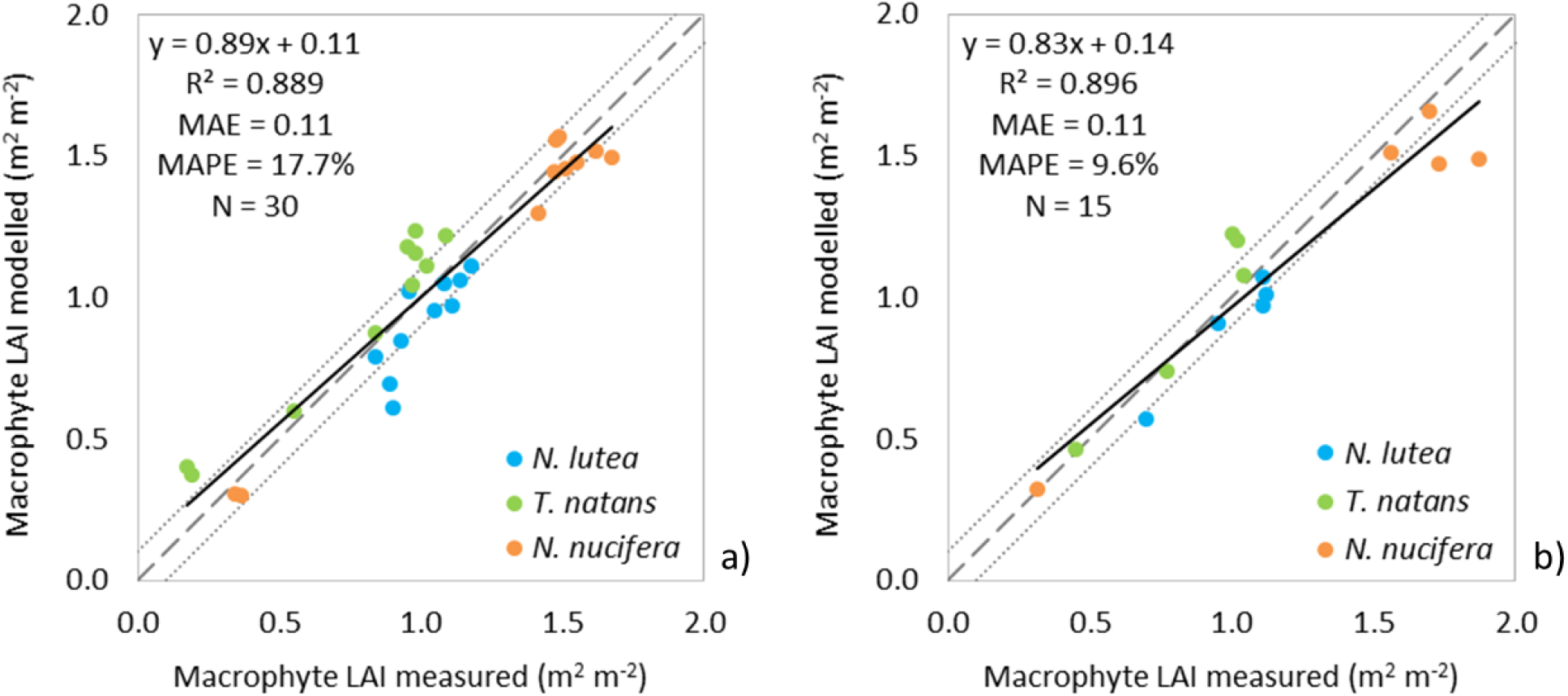
Comparison of macrophyte LAI measured in situ with estimates derived from the EVI2 based semi-empirical linear model implemented (SPOT5 scenes), over: a) calibration set; b) validation set. Dashed line is the 1:1, dotted lines mark the ±0.1 error line (N. *lutea* = *Nuphar lutea; T. natans* = *Trapa natans; N. nucifera* = *Nelumbo nucifera*).

### 5.2. Mapping macrophyte seasonal dynamics

#### 5.2.1. Mantua lakes system

Macrophyte seasonal dynamic maps derived for Mantua lakes system are reported in Figure 6. The time of the start of the season was quite different between the investigated lakes, in particular at Superior Lake. *N. nucifera* started to grow in mid-May (DOY 125-135), whereas *T. natans* appeared in mid-May (DOY 130-140) and early June (DOY 155-165) at Inferior and Middle lakes, respectively (Figure 8a). The peak of season was quite homogeneous (from late July to middle August) between species, except for *T. natans* at Inferior Lake where it was reached on mid-July (Figure 6b). In fact, for this latter stand the end of season was mid-late August, while the species disappeared at Middle Lake about one month later (Figure 6c). The last species that started the senescence period was *N. nucifera* (late October, Figure 6c).

**Figure 6.**
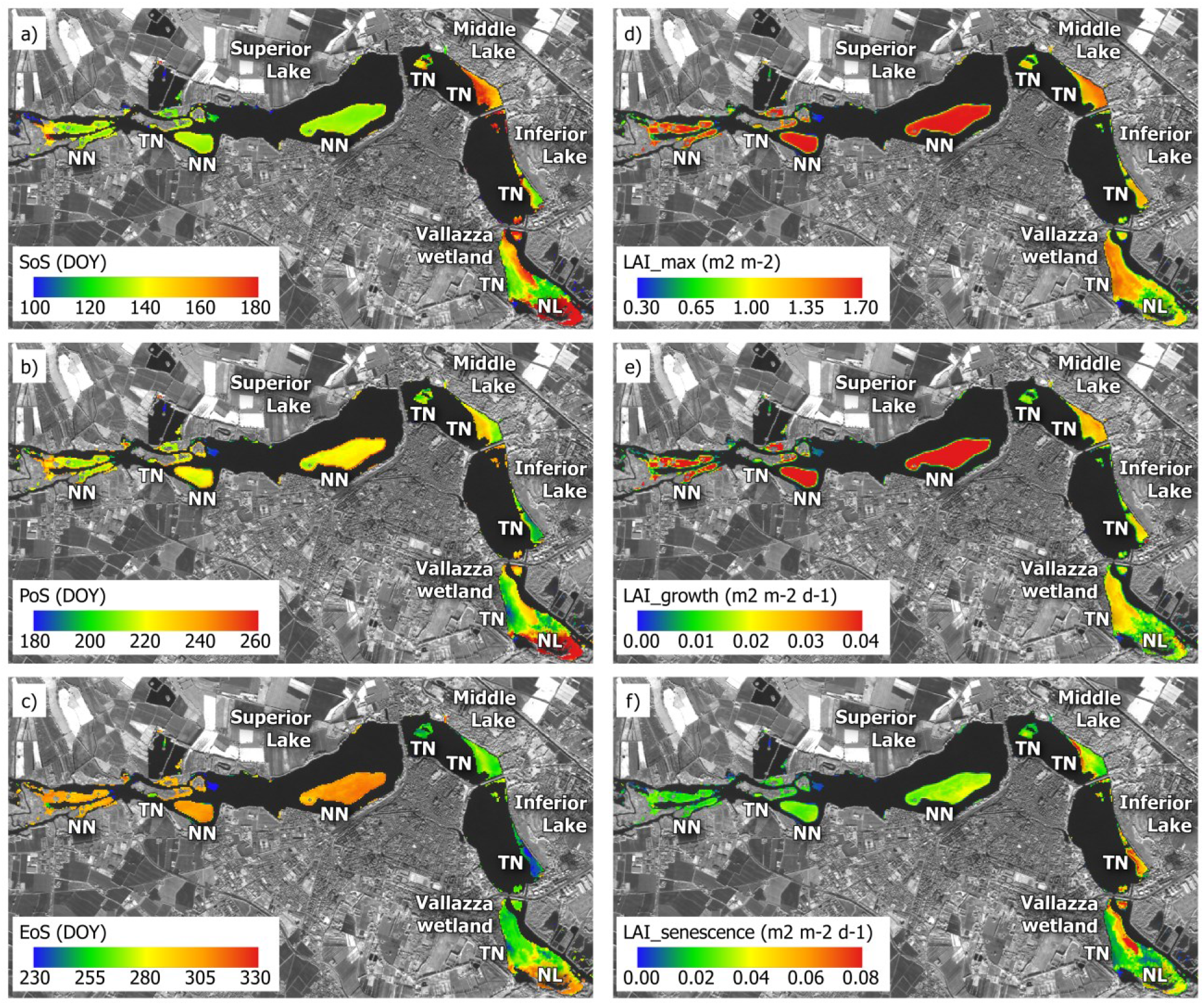
Macrophyte seasonal dynamics maps for Mantua lakes system study area: a) SoS; b) PoS; c) EoS; d) LAI_max; e) LAI_growth; f) LAI_senescence. SoS = start of season, PoS = peak of season, EoS = end of season, LAI_max = maximum LAI value, LAI_growth = rate of increase of LAI during the early growth, LAI_senescence = rate of decrease of LAI during the senescence. (NL = *Nuphar lutea;* TN = *Trapa natans;* NN = *Nelumbo nucifera*).

*N. nucifera* LAI values were higher than *T. natans* ones during the early and maximum vegetative phases; during the senescence phase the LAI decrease rates were more homogenous, especially at Middle Lake (Figures 6d, 6e, 6f). The length of season of *N. nucifera* and *T. natans* were up to 180 and 120 days, respectively. *T. natans* maximum LAI values at Middle Lake stand were higher than those at Inferior Lake (Figure 6d). In these two lakes the *T. natans* LAI values were similar during the early vegetative phase (Figure 6e), while during the senescence phase the LAI decrease rate was higher at Inferior Lake compared to Middle Lake one (Figure 6f). At Vallazza wetland, the macrophyte population was composed by different species, including *T. natans* and *N. lutea*, which reflects in more patched dynamics.

#### 5.2.2. Lac de Grand-Lieu

The seasonal dynamics maps derived for Lac de Grand-Lieu show a precocious development of the nymphaeids (*N. alba, N. lutea*) in the south of the central part of the lake, growing in protected bays with higher water transparency. The fastest species to develop is *T. natans*, which is highlighted by small blue spots in the middle of the lake in Figure 7b and 7c, showing anticipated PoS and EoS, around DOY 180 and 230, respectively. The peak value of LAI mapped for this species in 2015 (0.4-0.6) is lower than what observed in the field in previous years, when *T. natans* beds were more dense.

**Figure 7.**
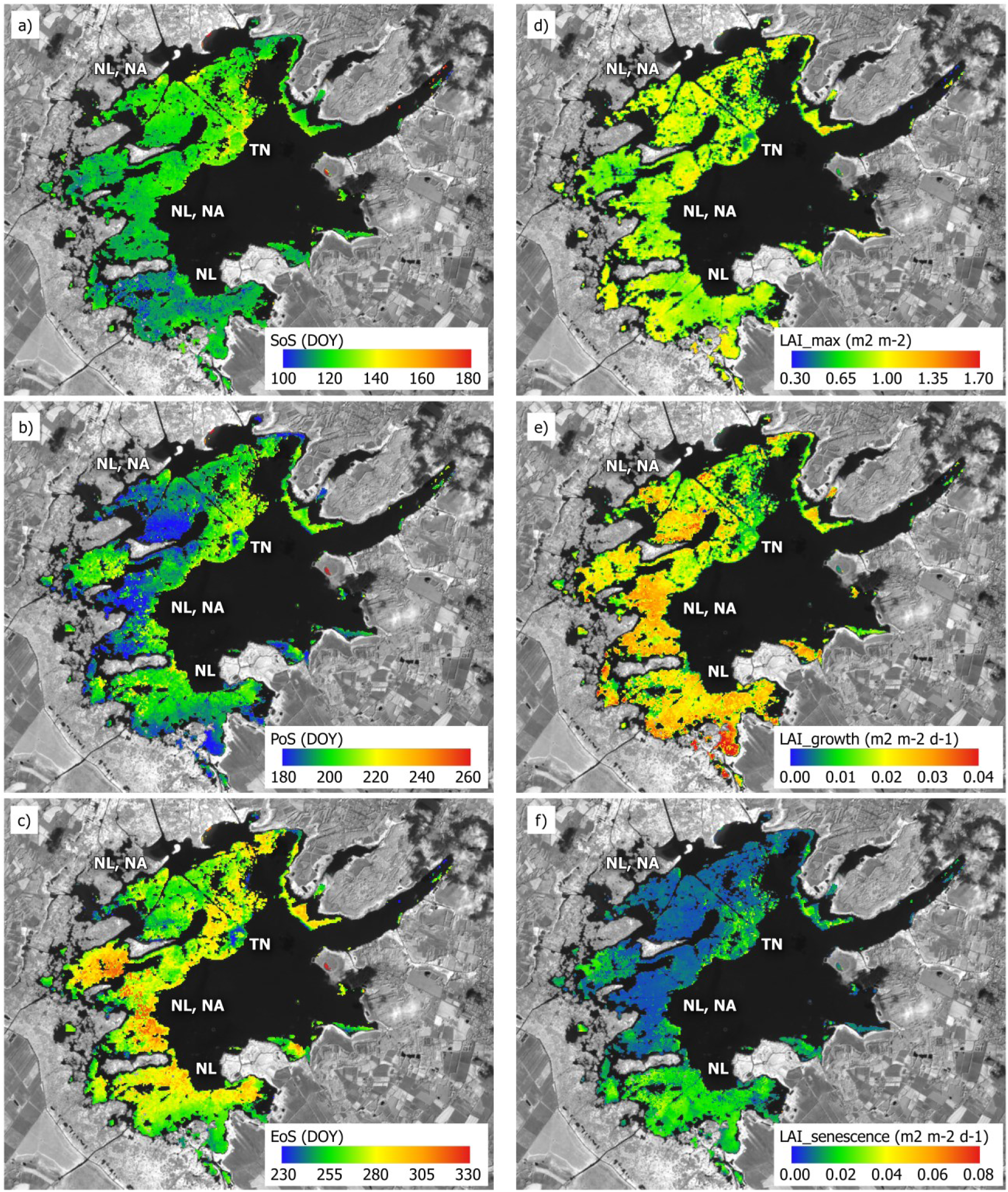
Macrophyte seasonal dynamics maps for Lac de Grand-Lieu test site: a) SoS; b) PoS; c) EoS; d) LAI_max; e) LAI_growth; f) LAI_senescence. Abbreviations are the same as in Figure 6. (NA = *Nymphaea alba;* NL = *Nuphar lutea;* TN = *Trapa natans*).

The early season development dynamics are shifted in time depending on the areas. The areas dominated by *N. lutea*, located in the south of the lake and with very low water level, show a precocious peak (DOY 180-185) and a high LAI growth rate. Additional analysis focusing on the balance between *N. alba* and *N. lutea* is needed in order to investigate the difference between the nymphaeids beds with similar SoS date and a moderate high LAI growth but a shifted vegetation peak. The areas with the latest EoS (DOY 295 to 330, Figure 7c) could be again attributed to dominance of *N. alba*, but the mixture of this species with *N. lutea* in most of the parts of the lake does not allow conclusive remarks without further data and analysis.

#### 5.2.3. Fundu Mare Island

The seasonal vegetation dynamics at Fundu Mare Island in 2015 reflects the changing hydrological conditions throughout the growing season. The coloured zones in Figure 8 mark the extent of the water covered area when the lakes are inundated completely (April-June, in 2015), before the lake water levels decreased in summer (Figure 3a).

**Figure 8.**
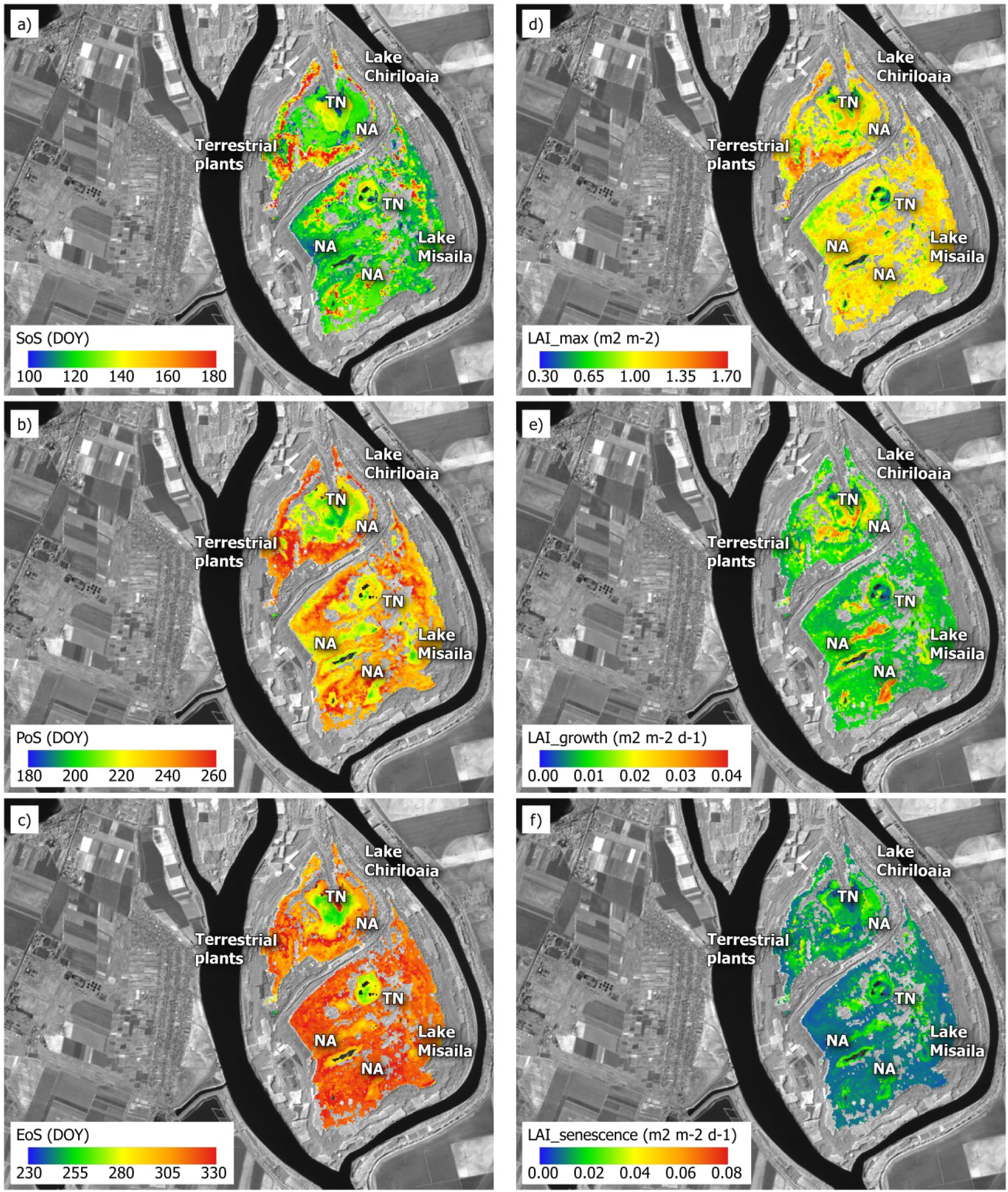
Macrophyte seasonal dynamics maps for Fundu Mare Island test site: a) SoS; b) PoS; c) EoS; d) LAI_max; e) LAI_growth; f) LAI_senescence. Abbreviations are the same as in Figure 6. (NA = *Nymphaea alba*; TN = *Trapa natans*).

The maps of macrophyte seasonal dynamics of Fund Mare Island show that the SoS for most of the area, in particular the aquatic ecosystems, occurred between DOY 116 and 132, i.e. end of April and beginning of May. SoS as late as on DOY 180 (late June) was observed in zones with massive willow encroachment. The PoS of floating plants ranges between 195 and 220 (mid to late July), and between 230 and 265 (August to September) for the zones that dried up when the water level dropped by around 2 m with respect to seasonal maximum (early August, Figure 3a). The EoS values show again a clear difference between the deepest lake areas with DOY 250 to 270 and outer zones that were dry during the later season.

The differences between the vegetation communities and the respective inundation duration are also reflected in the peak LAI value and its rate of change during growth and senescence. The highest peak LAI (1.35 to 1.65 m^2^ m^-2^) were mapped at the edges of Lake Chiriloaia (especially at North, West and South), where terrestrial species develop with low water level in late July. Small patches with LAI > 1.25 were found at many locations in both lakes and can be related to the willows-helophyte mosaic (see Figure 3b).

The highest rates of LAI change (Figures 8e and 8f) were observed for deeper lake areas, covered by floating and floating-leaved species, while the upland areas, inhabited by a mosaic of willows and helophytes, show more gradual growth and senescence.

### 5.3. Influence of input variables on macrophyte phenology estimation

#### 5.3.1. Influence of cloud cover amount

As shown in Table 5, under varying cloud cover threshold the absolute difference in phenology timing metrics with respect to the baseline (Mantua_5d dataset, see Table 4) is under 1.5 days, across all macrophyte species investigated. Averaged over all test sample beds, the MAD is lower than 0.8 days, and Max_span is under 1.9 days. Phenology timing outputs with maximum 50% cloud cover (Mantua_5d_CC50) show only slight differences with respect to the baseline (Max_span ≤ 0.34 days), while results using 30% and 10% maximum cloud cover are a little more diverging, scoring Max_span of 1.44 and 1.83 days, respectively. Among timing metrics tested, SoS is more sensitive to changes in cloud cover threshold choice (MAD ≤ 0.76 days), compared to PoS and EoS (MAD ≤ 0.31 days).

**Table 5.** Influence of cloud cover amount in input time series on TIMESAT output metrics (SoS, PoS, EoS) for Mantua lakes system dataset.

|  |  | Difference vs. baseline<br>(Mantua_5d) |  |  |
| --- | --- | --- | --- | --- |
| Dataset* | | ΔSoS<br>(days) | ΔPoS<br>(days) | ΔEoS<br>(days) |
| Mantua_5d_CC10 | NNs | 0.5 | -0.1 | -0.6 |
|  | TNm | 0.7 | -0.1 | 0.1 |
|  | TNi | -1.1 | 0.8 | 0.0 |
|  | NLv | -0.4 | 0.6 | 0.0 |
|  | **MAD** | **0.68** | **0.31** | **0.28** |
|  | *Max_span* | *1.83* | *0.93* | *0.76* |
| Mantua_5d_CC30 | NNs | -1.4 | 0.3 | 0.5 |
|  | TNm | 0.0 | 0.0 | 0.1 |
|  | TNi | -0.6 | 0.0 | -0.2 |
|  | NLv | -0.8 | -0.4 | 0.0 |
|  | **MAD** | **0.76** | **0.19** | **0.27** |
|  | *Max_span* | *1.44* | *0.75* | *0.70* |
| Mantua_5d_CC50 | NNs | 0.0 | 0.0 | 0.0 |
|  | TNm | 0.0 | 0.0 | 0.0 |
|  | TNi | 0.3 | 0.3 | -0.2 |
|  | NLv | 0.0 | 0.0 | 0.0 |
|  | **MAD** | **0.06** | **0.07** | **0.04** |
|  | *Max_span* | *0.25* | *0.34* | *0.17* |

Macrophyte test beds: NNs: *Nelumbo nucifera* (Superior Lake); TNm: *Trapa natans* (Middle Lake); TNi: *Trapa natans* (Inferior Lake); NLv: *Nuphar lutea* (Vallazza wetland).
\* see Table 4 for datasets description.

#### 5.3.2. Influence of temporal resolution

The differences in phenology timing metrics with respect to the baseline (Mantua_5d dataset, see Table 4) are summarized in Table 6, when the revisit of the input time series is reduced to 10 or 15 days.

**Table 6.** Influence of temporal resolution of input time series on TIMESAT output metrics (SoS, PoS, EoS) for Mantua lakes system dataset.

|  |  | Difference vs. baseline<br>(Mantua_5d) |  |  |
| --- | --- | --- | --- | --- |
| Dataset* | | ΔSoS<br>(days) | ΔPoS<br>(days) | ΔEoS<br>(days) |
| Mantua_10d_a | NNs | 1.8 | -3.8 | -3.6 |
|  | TNm | -0.3 | 0.7 | 2.4 |
|  | TNi | -0.7 | -0.4 | -1.1 |
|  | NLv | 1.3 | 1.4 | 0.2 |
| Mantua_10d_b | NNs | 0.9 | -0.4 | -0.5 |
|  | TNm | -2.6 | 0.5 | 0.8 |
|  | TNi | 0.3 | -0.1 | 2.2 |
|  | NLv | -0.8 | -1.0 | -0.5 |
| **Mantua_10d** | **MAD** | **1.19** | **1.15** | **1.59** |
|  | *Max-span* | *1.94* | *2.00* | *2.75* |
| Mantua_15d_a | NNs | 1.2 | 1.7 | 4.5 |
|  | TNm | -2.8 | 1.3 | 2.6 |
|  | TNi | -0.8 | -3.7 | -0.9 |
|  | NLv | 1.0 | 0.5 | 3.9 |
| Mantua_15d_b | NNs | -1.2 | -4.3 | -5.9 |
|  | TNm | -2.1 | -3.2 | -5.3 |
|  | TNi | 4.0 | 0.8 | -0.1 |
|  | NLv | 0.3 | -0.1 | -3.2 |
| Mantua_15d_c | NNs | 1.4 | -1.3 | -1.5 |
|  | TNm | -2.0 | -0.7 | -0.6 |
|  | TNi | -1.5 | -0.4 | 0.3 |
|  | NLv | -0.7 | 0.5 | -0.2 |
| **Mantua_15d** | **MAD** | **1.64** | **1.76** | **2.71** |
|  | *Max-span* | *3.09* | *4.46* | *7.35* |

Macrophyte test beds: NNs: *Nelumbo nucifera* (Superior Lake); TNm: *Trapa natans* (Middle Lake); TNi: *Trapa natans* (Inferior Lake); NLv: *Nuphar lutea* (Vallazza wetland).
\* see Table 4 for datasets description.

When the temporal resolution is degraded to 10 days, as for nominal Sentinel-2A coverage over Europe and Africa and for Sentinel-2A and −2B constellation globally, absolute differences in SoS, PoS and EoS across all macrophyte test beds, compared to the baseline, do not exceed 3.8 days. Different test beds, i.e. macrophyte communities, show variable sensitivity to reduction of time revisit at 10 days. The difference is generally comprised within 3.2 days (2 times the standard deviation across all test beds). Averaging over all macrophyte test beds, the MAD is lower than 1.6 days, and Max_span is under 2.8 days. Among timing metrics tested, EoS is more sensitive to reducing time revisit from 5 to 10 days (MAD = 1.59 days), compared to SoS and PoS (MAE ≤ 1.19 days).

Further reduction of temporal resolution of input time series up to 15 days, near to the nominal 16 day revisit of Landsat satellites (4 to 8) operationally collecting data all over the globe since 1982, results in absolute differences in TIMESAT output phenology metric up to 5.9 days across all macrophyte test beds. Influence on timing metrics is variable across macrophyte test beds investigated, and is generally comprised within 4.8 days difference to the baseline (2 times the standard deviation across all test beds). Averaging over all macrophyte test beds, the MAD of 15-day revisit time series can reach up to 2.71 days, while Max_span peaks at 7.35 days. As with 10-day reduced resolution, EoS is more sensitive (MAD = 2.71 days) than SoS and PoS (MAD ≤ 1.76 days).

#### 5.3.3. Influence of missing acquisitions

Based on timing metrics derived from the full Mantua lakes system dataset at 5-day temporal resolution (Figures 6a-c), five seasonal ranges were identified, namely: i) pre-season (DOY < 100); ii) early season, when most of macrophyte start of season dates occur (100 < DOY < 180); iii) full season, when peak of vegetative and reproductive phase for macrophyte communities investigated take place (180 < DOY < 250); iv) late season, after plant maturity is reached and senescence develops (250 < DOY < 320); and v) post-season (DOY > 320). These five seasonal ranges were used to interpret results of missing acquisitions influence.

At 5-day revisit (Figure 9a), EoS is the most sensitive parameter to missing a single date of the time series, with a MAD with respect Mantua_5d_CC50 dataset (see Table 4) reaching 3.5 days, while PoS and SoS are more robust to missing dates (MAD of 1.7 and 0.8, respectively). Sensitivity is pronounced for NNs test bed (populated by *N. nucifera*), with a maximum difference compared to the full dataset up to 8.3 days for EoS, and up to 4.2 days for PoS (i.e. when DOY 297 date is removed from the time series). The other macrophyte test samples, populated by *T. natans* and *N. lutea* communities, show lower sensitivity to missing acquisitions, with maximum difference to baseline lower than 1.5 days. Key periods for effectively capturing phenology timing from satellite LAI time series are therefore the early season (April-May) for SoS estimation, the late season (September-October) for PoS and EoS estimation, and late to post-season (October-November) for EoS estimation.

**Figure 9.**
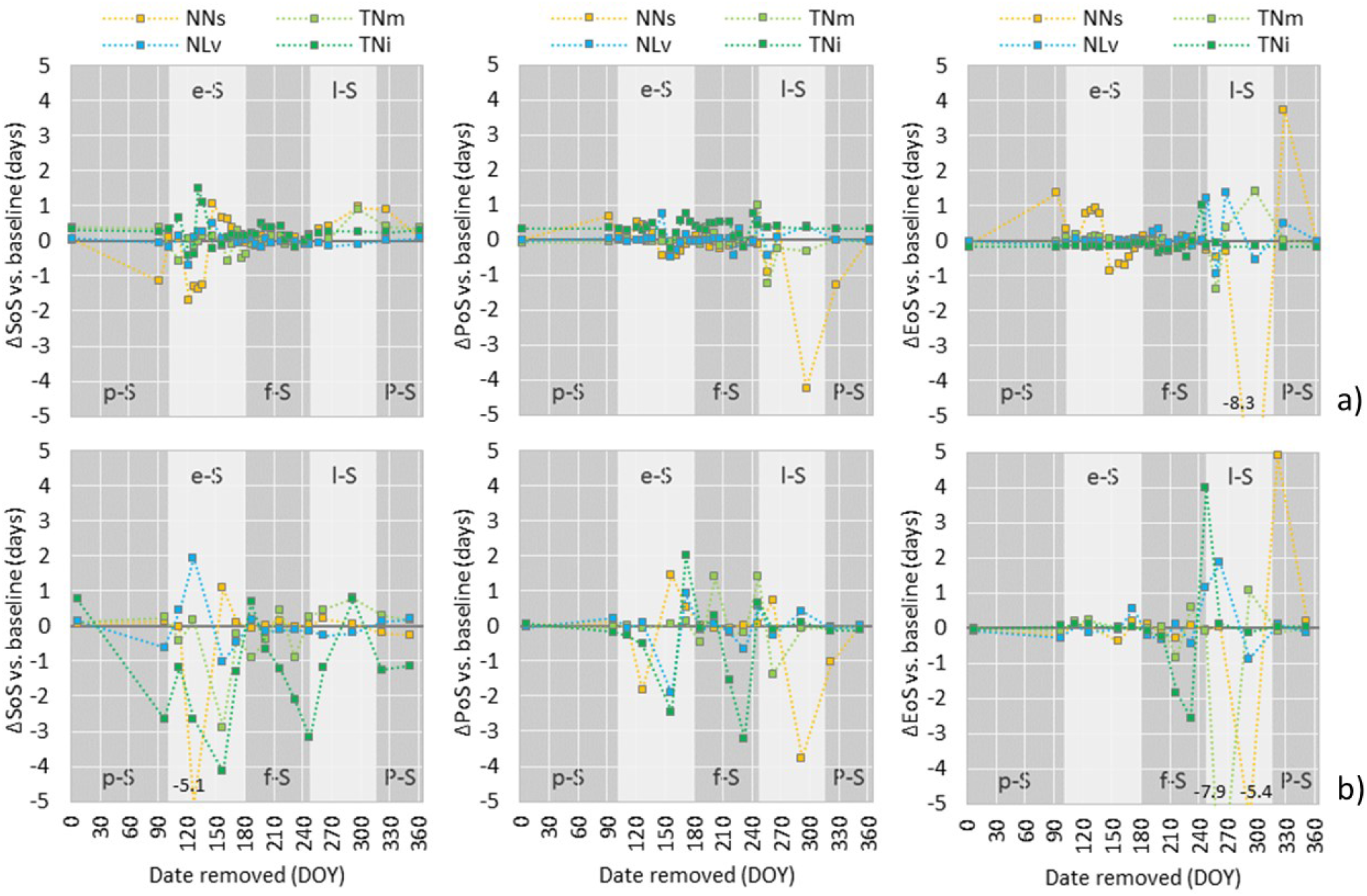
Influence of missing dates in the input time series on TIMESAT output metrics (SoS, PoS, EoS) for Mantua lakes system dataset, at: a) 5 day temporal resolution, and b) 15 day temporal resolution. Seasonal ranges highlighted by different grey colour background: p-S: preseason; e-S: early season; f-S: full season; l-S: late season; P-S: post-season. Macrophyte test beds: NNs: *Nelumbo nucifera* (Superior Lake); TNm: *Trapa natans* (Middle Lake); TNi: *Trapa natans* (Inferior Lake); NLv: *Nuphar lutea* (Vallazza wetland).

At 15-day revisit (Figure 9b), results are generally consistent with what highlighted for 5-day revisit, although with wider score ranges. SoS and EoS are the most sensitive parameters to missing a single date, with a MAD with respect to Mantua_15d_b dataset (see Table 4) around 2.7 days, while PoS is more robust (MAD of 1.5). Again, sensitivity is pronounced for NNs test bed, with a maximum difference compared to the full dataset up to 5.4 days for EoS, and up to 5.1 days for SoS. TNm test bed (populated by *T. natans*) shows high sensitivity to missing dates too, with differences up to 7.9 days for EoS, when dates in late to post-season time are removed from the time series. Sensitivity of timing metrics of TNi (also populated by *T. natans*) are less evident, and those for NLv (populated by *N. lutea*) are still the lower ones (< 2 days). At seasonal range level, key periods at 15-day temporal resolution are the same as for 5-day revisit.

## 6. Discussion

### 6.1. Macrophyte LAI mapping from satellite data

Satellite data coming from different medium resolution platforms (SPOT5, Landsat 7/8, Sentinel-2A) can be integrated into consistent time series of surface spectral reflectance with temporal misregistration under 0.3 pixels (Dai and Khorram, 1998) and radiometric mismatch across sensors contained within acceptable levels, i.e. a relative MAE < 0.04 between SPOT5 and Landsat 8 or Sentinel-2A data and an absolute MAE < 0.07 compared to *in situ* spectra (Figure S2). Compared to EVI2 derived from SPOT5 bands, both Landsat 8 and Sentinel-2A tend to score higher by around 30% in the high end of values (Figure S1). This is probably due to the difference in atmospheric correction algorithm used for the two types of data, MACCS for SPOT5 and ATCOR for Landsat and Sentinel-2, and implies that macrophyte LAI mapped may be slightly overestimated in the temporal range not covered by SPOT5 Take5 dataset. Nevertheless, being this range (before April and after September) mostly outside the growing season for macrophytes in temperate areas, the effect on TIMESAT derived phenology metrics is considered minor and therefore not significantly affecting the results of this study.

LAI for target macrophyte species (floating, floating-leaved, emergent) was reliably estimated using the semi-empirical regression model based on EVI2, as best performing spectral index: the MAE scored for independent validation data is 0.11 m^2^ m^-2^, with a tendency to underestimation due to index saturation only occurring for LAI > 1.5. These results demonstrate with experimental data covering an entire growing season that macrophyte LAI mapping based on spectral indices is feasible, with acceptable errors. The concept, already widely established in scientific literature for terrestrial vegetation, has been developed on a sound basis for aquatic vegetation only recently (see Villa et al., 2014), and this work confirms and put forward the findings of Villa et al. (2017), by extending the application potential macrophyte morphology mapping to broadband multispectral platforms.

The LAI model was calibrated (and validated) with reference *in situ* measures collected at the same spatial resolution (10 m grid) as the SPOT5 data. This is consistent with good practices and protocols (e.g. Weiss et al., 2004; Hufkens et al., 2008) delineated for the collection of vegetation biophysical characteristics to be linked to remotely sensed data at medium-high resolution (≤30 m pixel). Garrigues et al. (2006) reported that such practices are based on the assumption of homogeneous vegetation cover at this scale, which was verified by direct observation of macrophyte beds surveyed.

Within this work, we focused on floating and emergent growth forms of aquatic vegetation. Optical spectral response of submerged macrophytes is prevalently determined by the properties of water column above them, and using remotely sensed spectral reflectance for estimating their biophysical parameters is virtually not feasible in turbid systems (e.g. Hunter et al., 2010, Villa et al., 2015), as the majority of shallow lakes hosting abundant aquatic plant communities are. Even in case of coexistence between submerged and floating/emergent species and low turbidity, the high absorption of near-infrared light by clear water makes the influence of submerged macrophytes on the SIs tested and therefore on our estimates of macrophyte LAI practically negligible (e.g. Malthus, 2017).

### 6.2. Macrophyte seasonal dynamics at the study areas

Seasonal dynamics maps derived using TIMESAT with macrophyte LAI time series as input showed differences in both spatial and temporal patterns across the three study areas, temperate shallow ecosystems inhabited by common macrophyte species.

Nymphaeids, mainly *N. lutea* and *N. alba* in our study areas, show a growing season delayed by 10-30 days in Fundu Mare Island, compared to Mantua lakes system and Lac de Grand-Lieu, which is possibly an effect of water level fluctuation observed in the Romanian wetland (Figure 3a). Here, the LAI peak value for floating-leaved plants communities is higher than 1.1, due to interference of willow encroachment within *N. alba* beds (Figure 8d). In the Lac de Grand-Lieu, the seasonal dynamic of the nymphaeids can also affect the interpretation of the maps. *N. alba* can have a bimodal development with a first peak of biomass in spring and a second one late in summer or just one single peak of vegetation development (Paillisson and Marion, 2006). This dynamic depends on the years, and probably varies locally within the lake. Of all the sites, the growing season is the shortest and LAI growth rate the slowest in Mantua lakes system (Figure 6a-b and 6e), where nymphaeids cover a small area (< 6% of macrophyte surface).

Water chestnut (*T. natans*), the other species common to all three study areas, shows the highest intra-site variability of timing metrics (SoS, PoS, EoS) in Mantua lakes system, where different sub-systems link to spatially heterogeneous phenology. This shows the ability of *T. natans* to adapt to different environmental conditions and trophic levels, hence becoming dominant everywhere but in the Superior Lake and covering almost 50% of total macrophyte area.

On average, these species starts growing later in Mantua lakes system and reaches peak LAI values greater than 1.20 m^2^ m^-2^ (Figure 6d). Such high scores, anomalous for a floating plant, are due to the mixture with duckweed (lemnoids, such as *Lemna* ssp., *Spirodela polyrhiza*), coexisting with *T. natans* in Middle and Inferior lakes stands in full summer (July and August) and increasing the total leaf area per unit surface.

In the Lac de Grand-Lieu, where *T. natans* covers only a small area (< 5% of macrophyte surface) and is outcompeted by nymphaeids, growing season peaks and ends earlier than in the other two sites (Figure 7b-c), and the plant density reached is by far the lowest, with maximum LAI ~ 0.5-0.6 m^2^ m^-2^ (Figure 7d). The species has been in decline for the last forty years in Grand-Lieu, but the hydro-meteorological conditions in 2015 spring may have exacerbated the trend: a late flood at beginning of May raised lake level by 40 cm in 5 days, negatively affecting *T. natans* germination and growth.

The allochthonous sacred lotus (*N. nucifera*) is present only in Mantua lakes system, where it was introduced in 1921. The seasonal dynamics maps clearly show the distinct invasiveness traits of this species, characterized by fast growth (0.03-0.05 m^2^ m^-2^ d^-1^, from May to June; Figure 6e), long persistence (EoS in October-November; Figure 6c), high density and coverage (LAI up to 1.7 m^2^ m^-^; Figure 6d). These characteristics can easily explain how the autochthonous species (*T. natans* and *N. lutea*) were outcompeted by *N. nucifera* during the last century in Mantua lakes system, bringing to a situation where sacred lotus almost completely dominates Superior Lake, covering up to 45% of total macrophyte surface in the area, and needs to be managed by cutting on a yearly basis (Villa et al., 2017). This is a crucial point considering the very high number of allochthonous species spread in aquatic ecosystems, especially in temperate regions (Hussner, 2012; Bolpagni et al., 2013). They represent one of the most critical factors affecting the survival of autochthonous aquatic species and habitats (Gallardo et al., 2016).

In this study, the time for the start of the season was defined based on observation of target macrophyte species as the time when the macrophytes overpassed 25% of peak LAI. This may be different, however, in presence of diffused willow encroachment, as we found at Fundu Mare Island. Young *Salix* spp., partly inundated during the spring, developed only later in the season depending on the decreasing water levels, thus affecting the LAI mapped for macrophyte patches subject to encroachment and the interpretation of some seasonal dynamics maps (Figure 8). Riparian vegetation composition, structure and vigour responds rapidly to hydrological regime changes (e.g. Merritt et al., 2010; Johnson, 2000; Loheide and Booth, 2011). This study showed that seasonal dynamics metrics derived from TIMESAT, in particular PoS and EoS, could be related to specific vegetation community types. Vegetation zoning in floodplains, such as Fundu Mare Island, is closely related to the flooding duration and the mean summer water level (Ellenberg, 1996).

The use of medium resolution satellite data offers new opportunities for monitoring and studying on-going vegetation changes, for example the encroachment of willows in zones that were earlier mapped as aquatic habitats. Moreover, the description of vegetation seasonal dynamics in terms of LAI, which can be used for the parameterization of flow resistance of floodplain vegetation (Aberle and Järvelä, 2013), may support the application of hydrodynamic models for flow and sediment transport for complex riparian environments.

### 6.3. Influence of satellite data variables on phenology metrics

Assessing the impact of satellite data characteristics on macrophyte seasonal dynamics derived from TIMESAT brought to interesting results in terms of operational requirements as well as error levels one can expect when input time series is sub-optimal. Although exclusively based on empirical data covering Mantua lakes system, thanks to the dense revisit allowed by SPOT5 Take5 experiment (anticipating current operational features of Sentinel-2 constellation) our assessment provide a first, quantitative account of the sensitivity of key phenology timing metrics of aquatic vegetation to cloud cover of single dates, temporal resolution and missing acquisitions within input time series.

For 5-day revisit datasets, the effect of setting different cloud cover thresholds is contained under 2 days for the estimation of SoS, and even less for PoS and EoS. Temporal resolution is more decisive, reaching 2.8 and 7.4 maximum difference in EoS estimation, when time revisit of input time series is reduced from nominal to 10 and 15 days (similar to Landsat series revisit), respectively. The difference is less severe from SoS and PoS, for which the estimation bias can reach 1.8 and 3.1 days, respectively. Based on these results, Landsat data (4 to 8), acquired consistently from 1982 all over the globe, could be used for mapping phenology timing of macrophytes at 30 m spatial resolution in a retrospective way, with and error level around 2–3 days for SoS and PoS.

The analysis of influence of missing acquisitions, e.g. covered by clouds, highlights once again that EoS is the most sensitive timing parameter, bringing about estimation bias of 3.5 days on average, and 8.3 days at maximum, when satellite scenes are missing in the late season (from mid-September to mid-November). Differences for SoS and PoS are contained within 2 days and 5 days, respectively. Bias scores are higher for *N. nucifera*, while timing metric differences for *T. natans* and *N. lutea* are generally lower (< 1.5 days).

## 7. Conclusions

Our findings demonstrate that dense time series of different medium resolution satellite data (i.e. Landsat, SPOT, Sentinel-2) can be integrated to provide consistent maps of macrophyte LAI and their seasonal dynamics, although some criticality remain dealing with atmospheric correction using different algorithms and correspondence of TIMESAT derived metrics with actual phenological phases of different species. For the first time, the feasibility of reliably estimating macrophyte LAI with operational Earth Observation data was demonstrated, and the sensitivity of key phenology metrics was assessed with respect to cloud cover threshold used, temporal revisit and missing acquisitions in satellite time series used as input.

The use of satellite data for mapping macrophyte dynamics in a quantitative way at local to regional scales offers new possibilities for the monitoring of restoration and conservation actions in shallow aquatic ecosystems, i.e. the effect of measures affecting the hydrological conditions as well as the rapid changes that characterized the intra- and inter-seasonal macrophyte dynamicity. Furthermore, the results described confirm the effectiveness of remote sensing techniques in investigating driving factors of potential allochthonous species establishment, and in particular the competition of autochthonous vs. allochthonous species.

## Acknowledgements

This study has received funding from the European Community’s 7th Framework Programme, under project INFORM [grant no. 606865]. SPOT5 data have been acquired in the framework of SPOT5 (Take5) initiative, sponsored by CNES and ESA, and the time series covering Mantua lakes system was proposed within the MacroSentinel project [ESA project ID 29146].

The monitoring of the floating macrophytes in Lac de Grand-Lieu was supported by the Regional Directorate of Environment (DREAL) of Pays de la Loire, the regional Council of Pays de la Loire and Loire-Atlantique Federation of Hunters.

The investigations at Fundu Mare Island were funded by the European Economic Area (EEA) project ‘‘Restoration of the aquatic and terrestrial ecosystems of Fundu Mare Island” [grant no. RO02-0008] with financial contributions from Norway and Romania, and the support of the Natural Park Administration of the Small Wetland of Braila.

The authors thank Ilaria Cazzaniga (CNR-IREA) for her help during fieldwork in Mantua lakes system.

